# Hemato-Chemical Variations- A Diagnostic Tool for Babesia, Theileria and Anaplasma Infections in Captive Hog Deer (*Axis Porcinus*)

**DOI:** 10.1101/2024.04.23.588770

**Authors:** Muhammad Azhar, Muhammad Hassan Saleem, Ayesha Safdar, Muhammad Rizwan Khan, Bushra Nisar Khan, Rida Fatima, Furqan Awan, Muhammad Mujahid Khalid, Junaid Nadeem

**Author notes:** **Corresponding Author:** Professor Dr. Muhammad Hassan Saleem, Department of Veterinary Medicine, UVAS, Lahore, 00923334287286.

## Abstract

Hog deer are an essential component of Pakistan’s biodiversity. Their conservation helps maintain integrity of local ecosystems, contributing to overall stability and balance of region’s wildlife populations. In Pakistan, they are classified as an endangered species according to IUCN. Theileria, babesia and anaplasma are among most prevalent blood parasites in ruminants which require diagnosis at early stages of the infection so that treatment protocols can be followed and more severe pathological alterations can be prevented. The changes in hemato-chemical parameters caused by Theileria, babesia and anaplasma were recorded individually and then compared with healthy group of hog deer respectively. Alterations in hemato-chemical parameters produced by these parasite-infected hog deer were assessed as potential diagnostic tools for distinguishing between individual blood parasites. Total leukocyte count, monocytes and eosinophils were observed to be higher than normal in babesia infected hog deer while level was lower than normal hog deer in anaplasma and theileria infected hog deer. The main objective of this study is to promote early diagnosis of parasitic infections in hog deer to facilitate prompt treatment, pathological prevention and allocate resources for further research on hemato-chemical parameters as diagnostic tools, enhancing understanding and management of blood parasite infections in hog deer.

## INTRODUCTION

Theileria, babesia and anaplasma are amongst the general tick-borne protozoan that are spread around the world and infect nearly all warm-blooded animals in which they cause latent infection or in more severe cases, lethal infection [Moudgil and Singla, 2013; Novacco *et al*. 2019]. Apart from the risk of transmission to domestic animals, it is important to pay attention to the parasitic disorders which affect wild animals. It has frequently been claimed that wildlife serves as a reservoir for parasitic diseases that affect humans as well as animals [Choquette, 1956]. Understanding wildlife’s parasite biodiversity, how it serves as a reservoir for parasite infections, and how parasites are incorporated into the larger ecosystem are all extremely crucial factors when considering conservation of endangered and critically endangered species [Thompson *et al*. 2010].

Members of the Cervidae family play a significant role in a wide range of hemoparasites’ epidemiology. One of the significant elements affecting tick density is the presence of deers [Fanelli, 2021]. Theileria, a hemoprotozoan disease that is spread by ticks causes a cascade of symptoms which include hepatic and renal dysfunction, hypoproteinemia, hypoalbuminemia, and hypophosphatemia [Kannan, 2017]. *Anaplasma marginale* and *Anaplasma centrale* of the order Rickettsiales are the main causes of anaplasmosis, which was once referred to as gall sickness [Di Domenico *et al*. 2016; Sivajothi and Reddy, 2022]. Sheep, goats, and other ruminants in the wild are susceptible to *Anaplasma ovis* [Abd Ellah, 2015; Hornok *et al*. 2011].

Some forms of Babesia have been found in the family Cervidae [France, 2004]. The recurrence of the disease may be brought on by stressors such as inadequate nourishment, the rutting season, and travelling [Fanelli, 2021]. Ticks, wild ruminants, and infected livestock can all act as A. marginale reservoirs [De La Fuente *et al*. 2004].

A drop in the concentration of hemoglobin, PCV, and red blood cell counts under high parasitemia are typically indicative of clinical babesiosis. An increase in total leukocyte count and thrombocytopenia are specific hematological characteristics of babesia infection [Novacco *et al*. 2019]. While microcytic hypochromic anemia, leukocytopenia, significant thrombocytopenia, increased lymphocytes, and leucopenia are some of the common findings in the cases of theileriosis [Abd Ellah, 2015]. Lastly, hematological analysis of anaplasma positive animals reveals decreased hemoglobin levels, a rise in leucocyte count, a total erythrocytic count, and PCV, elevated levels of ALT, BUN, bilirubin, and alkaline phosphatase [Sivajothi and Reddy, 2022]. Leukocytosis observed in parasitic infections is an indication towards the fact that lymphoid organs have been stimulated by parasites and their poisons [Kuttler, 1984].

A fundamental problem in the protection of threatened species is the impact that parasites have on the dynamics of wildlife populations. It is evident that gathering and analyzing information about the host/parasite relationships of parasites plays a significant role in the conservation of various wildlife species [Malan *et al*. 1997; Sanchez *et al*. 2018].

In the following study, various hemato-chemical parameters were assessed to determine the variations caused by the hemoparasites; theileria, babesia and anaplasma in captive hog deer from around Pakistan. The findings of the investigations undertaken used to establish the baseline data on hemato-chemical variations individually for Babesia, Theileria and Anaplasma and then compared with each other for differential diagnosis. Ultimately, this research is a step towards enhancing our knowledge of the interactions between tick-borne pathogens and deer populations, helping to inform more effective wildlife management and conservation strategies in regions where these infections are prevalent.

## MATERIALS AND METHODS

### SAMPLE COLLECTION

Aseptic blood samples up to 6 ml for each animal were taken using 18-gauge disposal syringes from the jugular veins. Blood was drawn 3ml placed in an EDTA vial at a concentration of 2 mg/ml as an anticoagulant and the other 3ml in non-EDTA vial. For this study, a sample size of n=4 PCR-positive deer were selected for each of the three conditions: theileria, babesia, and anaplasma.

### ASSESSMENT OF HEMATOLOGICAL PARAMETERS

The hematological parameters under investigation included a complete blood count (CBC), encompassing hemoglobin (Hb), packed cell volume (PCV), total leukocyte count (TLC), total erythrocyte count (TEC), mean corpuscular volume (MCV), mean corpuscular hemoglobin (MCH), mean corpuscular hemoglobin concentration (MCHC), as well as levels of neutrophils, eosinophils, lymphocytes, basophils, and monocytes.

### ASSESSMENT OF BIOCHEMICAL PARAMETERS

Additionally, liver function tests (LFTs) were conducted, which included the evaluation of aspartate aminotransferase (AST) and alanine aminotransferase (ALT) levels, while renal function tests (RFTs) included assessments of creatinine and urea levels. Calcium, phosphorus, sodium, and potassium levels were also analyzed to provide a comprehensive overview of the hemato-chemical alterations associated with these infections in captive hog deer (Mitema *et al*. 1991).

### MOLECULAR DIAGNOSIS

After preliminary screening of suspected animals through clinical signs and simple microscopy, the microscopic positive samples were subjected to PCR for confirmatory diagnosis. The details of molecular diagnosis of each blood parasite are as under;

In a Polymerase Chain Reaction (PCR), the initial denaturation; 95 ºC for 1 minute. The cyclic denaturation phase begins; 95 ºC for 30 seconds in each cycle. Next is the annealing phase; 60 ºC for 45 seconds during each cycle. The extension; 72 ºC for 30 seconds. There is a then the final extension at 72 ºC for 10 minutes. For the amplification of DNA associated with Babesia, the forward primer; F1; 5’-GTCTTGTAATTGGAATGATGG-3’. The reverse primer; R1; 5’-TAGTTTATGGTTAGGACTACG-3’. A total length DNA fragment of 465 base pairs was produced [Casati *et al*. 2006].

The forward primer; 5′-TAGTGACAAGAAATAACAATACGGGGCTT-3′, and the reverse primer; 5’-CAGCAGAAATTCAACTACGAGCTTTTTAACT-3′, were crafted to target the DNA region associated with Theileria. A DNA fragment measuring 149 base pairs in length was produced (Yang *et al*. 2014). The forward primer; MSP3F; 5′-CCA GCG TTT AGC AAG ATA AGA G-3′, and the reverse primer; MSP3R; 5′-GCC CAG TAA CAT CAT AAG C-3′. These primers were chosen to target a specific DNA region associated with Anaplasma. A DNA fragment measuring 334 base pairs in length was produced [Kim *et al*. 2017].

### ASSESSMENT OF NORMAL HEMATO-CHEMICAL PARAMETER VALUES

For this purpose, n=04 apparently healthy, PCR negative and Pakistani origin captive hog deer 10 (Attributed as Control group) were selected and used for the assessment of normal hemato-chemical parameter values to be compared to the collected PCR positive samples. These values were used as reference values for comparison with diseased hog deer.

### STATISTICAL ANALYSIS

The data on hemato-chemical variations was analyzed through simple T-test by using R software. P<0.05 at 95% CI was considered significant.

## RESULTS

In the comparative analysis of hemato-chemical parameters observed amongst the Theileria, Babesia, Anaplasma-positive hog deer (n=4 for each parasite) and the control group, notable distinctions were observed. For Theileria-positive hog deer group, the following parameters exhibited values lower than those of the control group, indicative of deviations from the normal (control) values: Hb, PCV, TEC, TLC, MCV, MCHC, Neutrophils, Lymphocytes, Monocytes, Basophils, Platelets, Calcium, Phosphorus, Sodium, and Potassium. Moreover, the Total Protein level was notably very low in this group. Conversely, the values of Total Bilirubin, ALT, BUN, and Creatinine exhibited higher values compared to the normal (control) values, signifying an elevation in these parameters in the Theileria-positive hog deer group. Remarkably, the AST value in the same group was substantially elevated in comparison to the control group as shown in the Table [1].

**Table 1:**
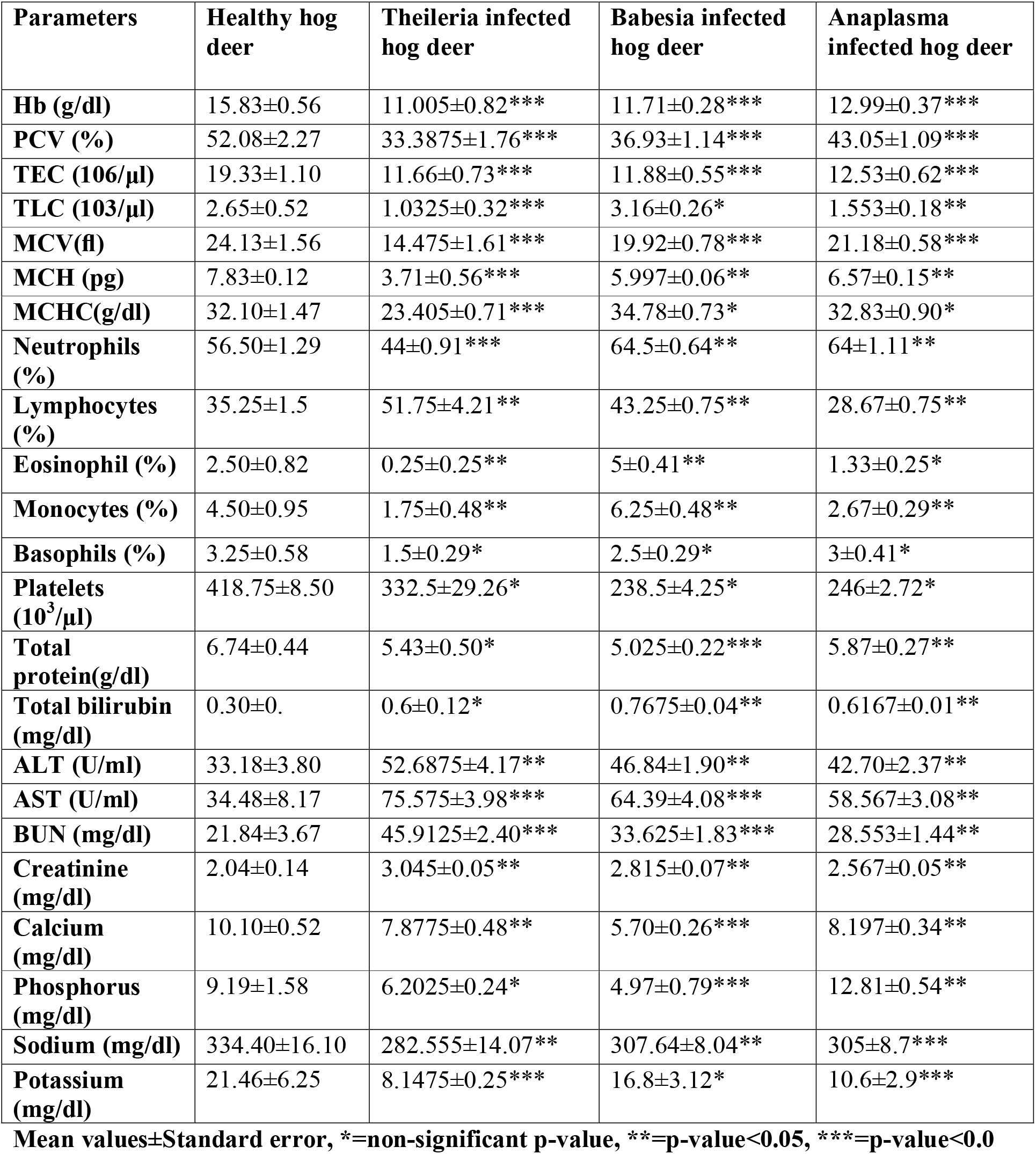
Hemato-chemical parameters of healthy hog dear observed amongst the Theileria, Babesia and Anaplasma.

| Parameters | Healthy hog deer | Theileria infected hog deer | Babesia infected hog deer | Anaplasma infected hog deer |
| --- | --- | --- | --- | --- |
| **Hb (g/dl)** | 15.83±0.56 | 11.005±0.82*** | 11.71±0.28*** | 12.99±0.37*** |
| **PCV (%)** | 52.08±2.27 | 33.3875±1.76*** | 36.93±1.14*** | 43.05±1.09*** |
| **TEC (106/µl)** | 19.33±1.10 | 11.66±0.73*** | 11.88±0.55*** | 12.53±0.62*** |
| **TLC (103/µl)** | 2.65±0.52 | 1.0325±0.32*** | 3.16±0.26* | 1.553±0.18** |
| **MCV(fl)** | 24.13±1.56 | 14.475±1.61*** | 19.92±0.78*** | 21.18±0.58*** |
| **MCH (pg)** | 7.83±0.12 | 3.71±0.56*** | 5.997±0.06** | 6.57±0.15** |
| **MCHC(g/dl)** | 32.10±1.47 | 23.405±0.71*** | 34.78±0.73* | 32.83±0.90* |
| **Neutrophils (%)** | 56.50±1.29 | 44±0.91*** | 64.5±0.64** | 64±1.11** |
| **Lymphocytes (%)** | 35.25±1.5 | 51.75±4.21** | 43.25±0.75** | 28.67±0.75** |
| **Eosinophil (%)** | 2.50±0.82 | 0.25±0.25** | 5±0.41** | 1.33±0.25* |
| **Monocytes (%)** | 4.50±0.95 | 1.75±0.48** | 6.25±0.48** | 2.67±0.29** |
| **Basophils (%)** | 3.25±0.58 | 1.5±0.29* | 2.5±0.29* | 3±0.41* |
| **Platelets (10<sup>3</sup>/µl)** | 418.75±8.50 | 332.5±29.26* | 238.5±4.25* | 246±2.72* |
| **Total protein(g/dl)** | 6.74±0.44 | 5.43±0.50* | 5.025±0.22*** | 5.87±0.27** |
| **Total bilirubin (mg/dl)** | 0.30±0. | 0.6±0.12* | 0.7675±0.04** | 0.6167±0.01** |
| **ALT (U/ml)** | 33.18±3.80 | 52.6875±4.17** | 46.84±1.90** | 42.70±2.37** |
| **AST (U/ml)** | 34.48±8.17 | 75.575±3.98*** | 64.39±4.08*** | 58.567±3.08** |
| **BUN (mg/dl)** | 21.84±3.67 | 45.9125±2.40*** | 33.625±1.83*** | 28.553±1.44** |
| **Creatinine (mg/dl)** | 2.04±0.14 | 3.045±0.05** | 2.815±0.07** | 2.567±0.05** |
| **Calcium (mg/dl)** | 10.10±0.52 | 7.8775±0.48** | 5.70±0.26*** | 8.197±0.34** |
| **Phosphorus (mg/dl)** | 9.19±1.58 | 6.2025±0.24* | 4.97±0.79*** | 12.81±0.54** |
| **Sodium (mg/dl)** | 334.40±16.10 | 282.555±14.07** | 307.64±8.04** | 305±8.7*** |
| **Potassium (mg/dl)** | 21.46±6.25 | 8.1475±0.25*** | 16.8±3.12* | 10.6±2.9*** |

Mean values±Standard error, \*=non-significant p-value, \*\*=p-value<0.05, \*\*\*=p-value<0.0

Specifically, significant changes were noted in the p-values of Hb, PCV, TEC, TLC, MV, MCH, MCHC, Neutrophils, Lymphocytes, Monocytes, Total Protein, AST, ALT, Creatinine, BUN, Calcium, Sodium, and Potassium. These alterations signify significant deviations from the baseline values. In contrast, the parameters of Basophils, Platelets, Total Bilirubin, and Phosphorus exhibited p-values that did not show significant alterations, indicating relative stability in these parameters under study as demonstrated in the Table [1].

Among the hematological parameters assessed and compared from a cohort of babesia-positive hog deer and a control group, the values of Hb, PCV, TEC, MCV, MCH, Basophils, Total Protein Level, Calcium, Phosphorus, Sodium, and Potassium exhibited a notable decrease relative to the control group. Furthermore, the platelet count in babesia-positive hog deer was found to be significantly reduced. Conversely, an elevation in the levels of TLC, MCHC, Neutrophils, Lymphocytes, Eosinophils, Monocytes, Total Bilirubin, ALT, BUN, and Creatinine was observed when compared to the control group. Particularly noticeable was the markedly elevated AST level in babesia-positive hog deer, indicating potential hepatic stress or damage. The results can be seen in the Table [1].

Significant alterations were associated with the following hematological and biochemical parameters in the cohort under investigation: Hb, PCV, TEC, MCV, MCH, Neutrophils, Lymphocytes, Eosinophils, Monocytes, Total Protein, Total Bilirubin, ALT, AST, BUN, Creatinine, Calcium, Phosphorus, and Sodium. Conversely, no significant alterations were observed in the p-values associated with the following parameters: TLC, MCHC, Basophils, Platelets, and Potassium.

In the comparative analysis of hematological parameters between Anaplasma-positive hog deer and a control group, it was observed that the Anaplasma-positive group exhibited significantly altered values in various hematological parameters when compared to the control group. The levels of Hb, PCV, TEC, TLC, MCV, MCH, lymphocytes, eosinophils, monocytes, basophils, total protein, and calcium were notably lower than the corresponding normal levels observed in the control groups. Additionally, platelet counts were significantly reduced in the anaplasma-positive group.

Conversely, neutrophil counts, as well as levels of ALT, AST, BUN, and phosphorus, were substantially elevated in the Anaplasma-positive group compared to the control group. Meanwhile, MCHC and creatinine levels exhibited only slight increases relative to the control group.

Levels of Hb, PCV, TEC, TLC, MCV, MCH, neutrophils, lymphocytes, monocytes, total protein, total bilirubin, ALT, AST, BUN, creatinine, calcium, and phosphorus demonstrated statistically significant changes as depicted in the Table [1]. Conversely, the values of MCHC, eosinophils, basophils, and platelets did not exhibit statistically significant alterations in the Anaplasma-positive group when compared to the control group.

### COMPARATIVE STUDY/ DIFFERENTIAL DIAGNOSIS

In the current study, the values of Hb, PCV and TEC are observed to be higher in anaplasma positive hog deer compared to the theileria and babesia positive hog deer. The value of TLC, eosinophils and monocytes appears to be higher in the babesia positive hog deer compared to the anaplasma and Theileria-positive groups as shown in the Figure [1]. The value of MCHC, and basophils is lower in theileria positive hog deer compared to the other two parasitic infestations. Lastly, lymphocytes levels were observed to be are lower than the normal value in anaplasma positive hog deer and higher than the values for the control group in the babesia and theileria positive hog deer groups.

**Figure 1:**
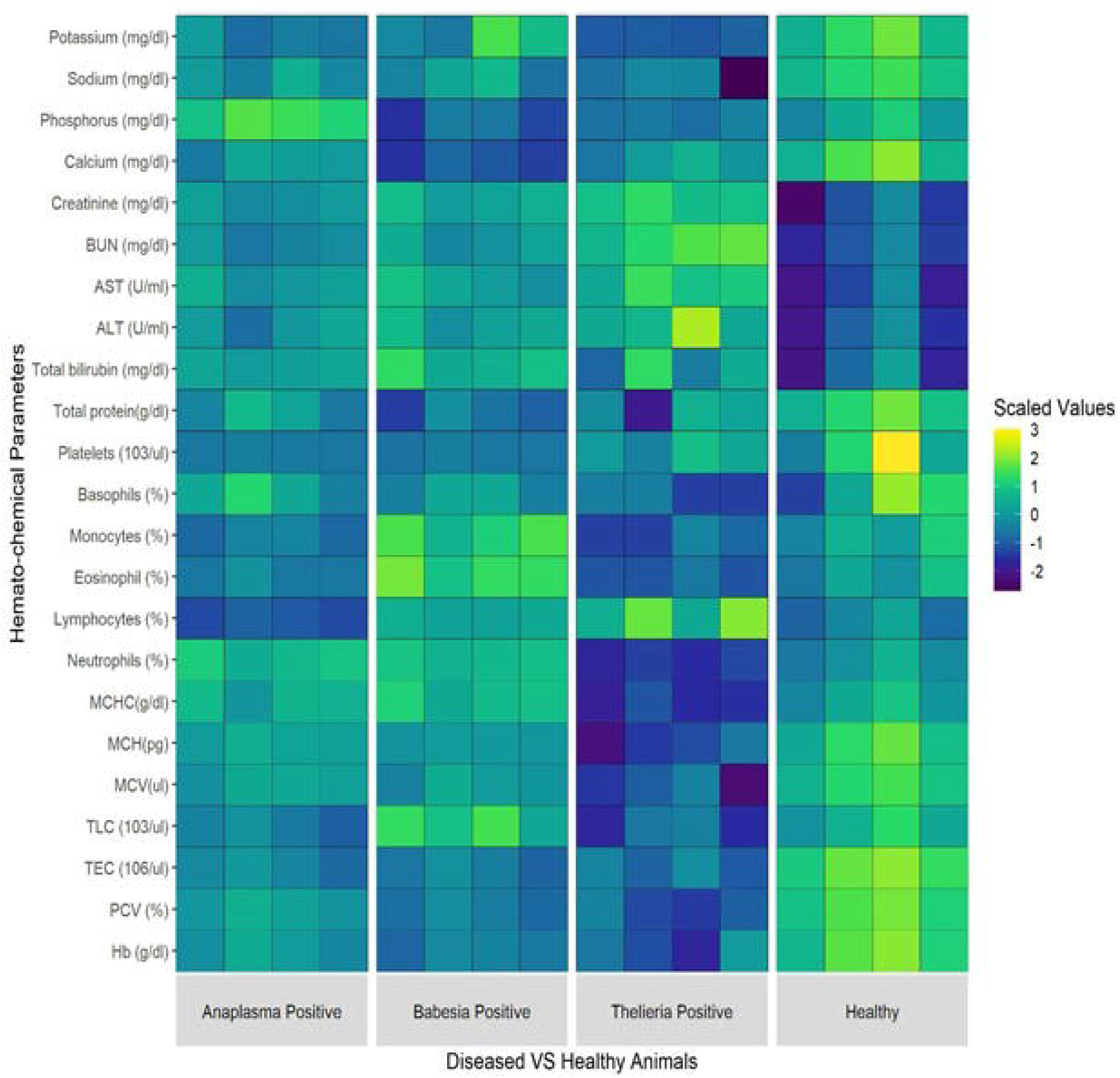
The comparative hemato-chemical variations in scaled values in diseased (Babesia, Theileria and Anaplasma infected) and healthy (non-infected) captive hog deer groups.

## DISCUSSION

The significance of blood parasites is due to the severe economic losses and their effect on the immune status of the body of the target animals [Alvarado-Rybak *et al*. 2016; Hussein *et al*. 2007]. Early diagnosis allows for prompt intervention and treatment of the otherwise deadly parasitic infections [Diaz *et al*. 2021; Thompson *et al*. 2009].

An outstanding feature of erythrocytic infections is the occurrence of anaemia. Anemia in anaplasmosis results from extravascular destruction of erythrocytes infested by parasites and extensive erythrophagocytosis initiated by parasitic damage to erythrocytes [Ahmadi-hamedani *et al*. 2012; Ajayi *et al*. 1987].

In most anaemic conditions, alteration in average size of red cells is often preceded by similar changes in MCH and often the MCHC. With microcytic cells, the MCH is decreased and this is referred as a microcytic hypochromic anaemia in anaplasma while normocytic hemolytic anemia is observed in theileria and babesia. Similar findings have been reported by scientists in the past [Chauvin et al. 2004]. Due to hemolytic anemia, the animal cannot tolerate stress, leading to fatigue and weakness [Borgsteede, 1996; Sivajothi and Reddy, 2022].

Hematological findings during clinical babesiosis are normally characterized by a decrease in hemoglobin concentration, PCV and red blood cell counts during high parasitemia, while total leukocyte count may increase [Novacco *et al*. 2019]. In the current study, Hemoglobin, PCV, and erythrocytes were observed to be lower than normal in the parasite infected hog deer. All three parasite infestations displayed a significant decrease in peripheral leukocytes, with the exception of babesia, where eosinophils and monocytes were found to be higher than the normal leucocyte levels. The decrease in the leukocyte levels is attributable to a severe and transient B-cell lymphopenia, followed by an intense and enduring neutropenia and eosinopenia [Ahmadi-hamedani *et al*. 2012].

The insignificant changes in TLC level in the babesia-infected group may be due to the destruction of red blood cells by the protozoan stimulating phagocytic cells such as lymphocytes and monocytes to clean up the toxic remnants of ruptured red blood cell [Chauvin *et al*. 2004]. Leukocytosis indicates stimulation of lymphoid organs and systems due to parasites and their toxins [Sivajothi and Reddy, 2022]. Contrarily, the decrease in the TLC in the anaplasma and Theileria-positive hog deer might be attributed to persistent harmful effects of toxic metabolites of Theileria and anaplasma on the haemopoietic organs especially bone marrow and their interference with the process of leucogenesis [Hussein *et al*. 2007; Woldehiwet, 2006].

Lower PCV and higher AST, and ALT indicate liver dysfunction [Sivajothi and Reddy, 2022]. On the progress of the severity of parasitic infections, there is an increase in the level of BUN as well as the total bilirubin. Significant increase in the activity of AST was also observed which indicates the harmful effect of toxic metabolites of babesia, theileria and anaplasma on the organs like soft tissue, liver, kidney. These results are supported by the findings of others [Chauvin *et al*. 2004; Dugat *et al*. 2014]. No significant difference was recorded in creatinine levels in animals found positive for Theileria and babesia according to previous studies [Ganguly *et al*. 2019; Hilpertshauser *et al*. 2006]. However, in the present study, the level of creatinine was observed to be significantly higher than normal in all three parasite infected hog deer groups indicating compromised kidney function as shown in the Figure [1].

The concentration of BUN in diseased animals may be due to increased turnover of proteins [Bishop *et al*. 2004; Sivajothi and Reddy, 2022]. The level of BUN in the present study was observed to be markedly higher in theileria positive hog deer as compared to babesia and anaplasma infected hog deer. Significant increase in bilirubin levels in affected animals may be due to hemolytic crisis of blood parasites [Chauvin et al. 2004]. Hyperbilirubinemia in all the parasite infected hog deer is due to excessive destruction of erythrocytes and indirect hepatocellular damage. During the disease process, the animals’ hemopoietic system is activated in response to erythrophagocytosis. The damage to the skeletal or heart muscles, hepatic tissues, and erythrocytes may result in a considerable increase in the level of AST and ALT [Cole and Friend, 1999; Sivajothi and Reddy, 2022].

## CONCLUSION

The conducted research on hematological variations in hog deer has provided valuable insights into diagnostic parameters for this species. Analysis of hematological and biochemical values, as explored in studies on other deer species offers a foundation for understanding the health status of hog deer. The changes observed in blood parameters of hog deer infected with babesia, theileria and anaplasma as demonstrated in the study provide crucial data for diagnosing potential parasitic infestations in hog deer. These findings contribute to the broader knowledge of hematological variations in different deer species and offer a basis for establishing species-specific diagnostic criteria.

## RECOMMENDATIONS

In the future, extensive studies should be conducted on hog deer specifically, exploring a wider range of haematological and biochemical parameters to enhance diagnostic precision. Additionally, the findings from the broader researches on other deer species need to be constantly compared to identify the common trends and unique characteristics in hog deer

