## Supplemental Data 1 for "Hemato-Chemical Variations- A Diagnostic Tool for Babesia, Theileria and Anaplasma Infections in Captive Hog Deer (*Axis Porcinus*)"

**Ethical/ IRB approval**

No animals were harmed throughout the study, which was conducted in accordance with Institutional Review Committee vide letter no. DR/402 dated 30 September 2021 issued from University of Veterinary and Animal Sciences Lahore, Pakistan.

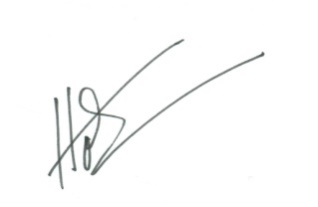

**Corresponding author**:

Dr. Muhammad Hassan Saleem

Professor

Department of Veterinary Medicine

University of Veterinary and Animal Sciences, Lahore, Pakistan

Contact# +923334287286
